## Supplementary material for "Targeting SLC7A11-mediated cysteine metabolism for the treatment of trastuzumab resistant HER2 positive breast cancer": Figures S1-S15 and Tables S1-S3

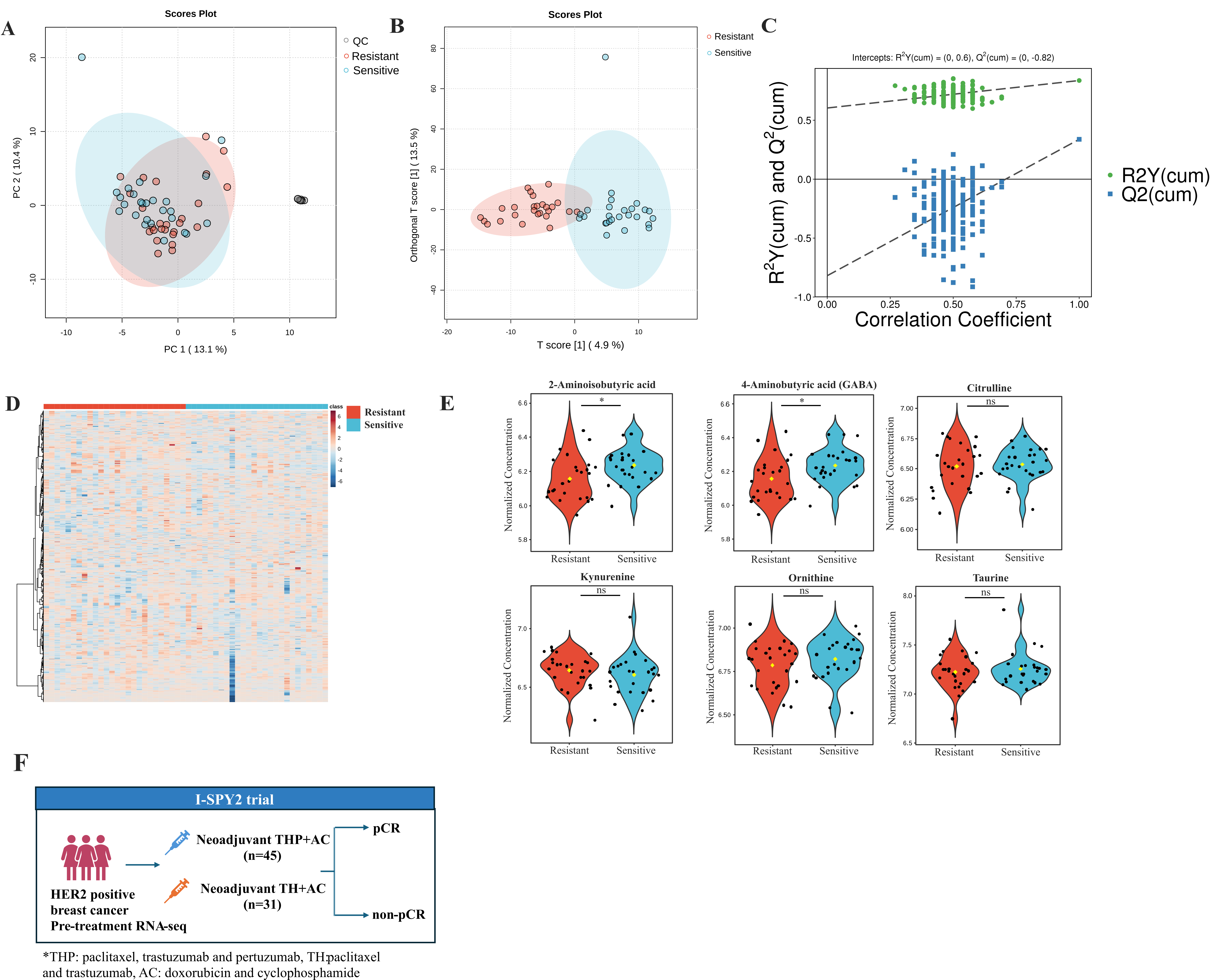

**Figure S1.** Quality control and differentiation analysis of plasma metabolites in trastuzumab sensitive and primary resistant HER2 positive breast cancer patients. (A) The distribution of quality control samples in PCA plots. (B) OPLS-DA of plasma metabolites in trastuzumab sensitive and primary resistant patients. (C) The robustness of the OPLS-DA model. (D) Heatmap of plasma metabolites in trastuzumab sensitive and primary resistant patients. (E) Violin plots of different non-protein amino acids in trastuzumab sensitive and primary resistant patients. (F) Overview of the I-SPY2 trial in transcriptomic analysis.

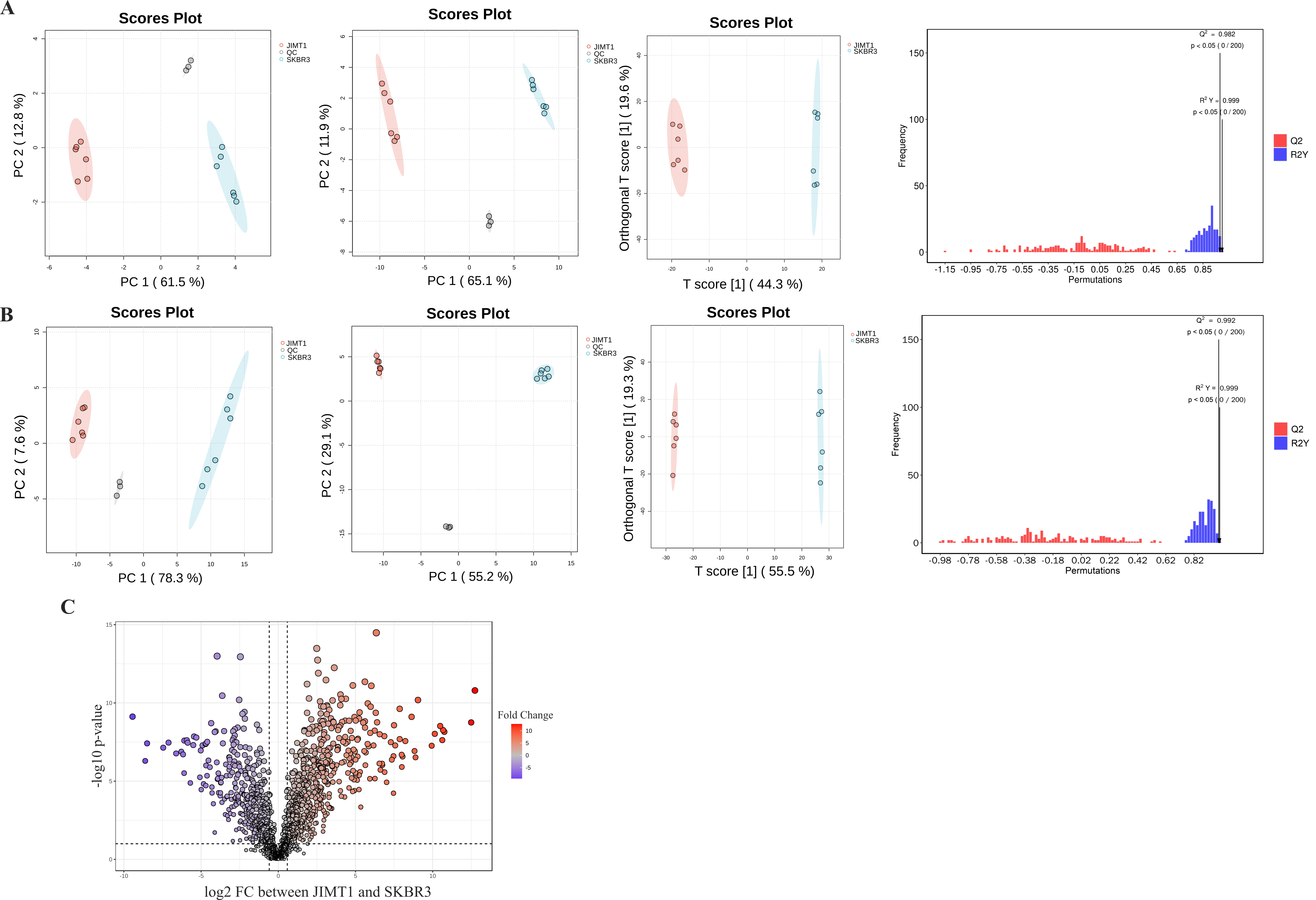

**Figure S2**. Quality control and differentiation analysis of metabolomic data in JIMT1 and SKBR3. (A) Principal component analysis (PCA) and orthogonal partial least squares-discriminant analysis (OPLS-DA) of hydrophilic metabolites. (B) PCA and OPLS-DA of lipophilic metabolites. (C) Volcano plot of different lipophilic metabolites.

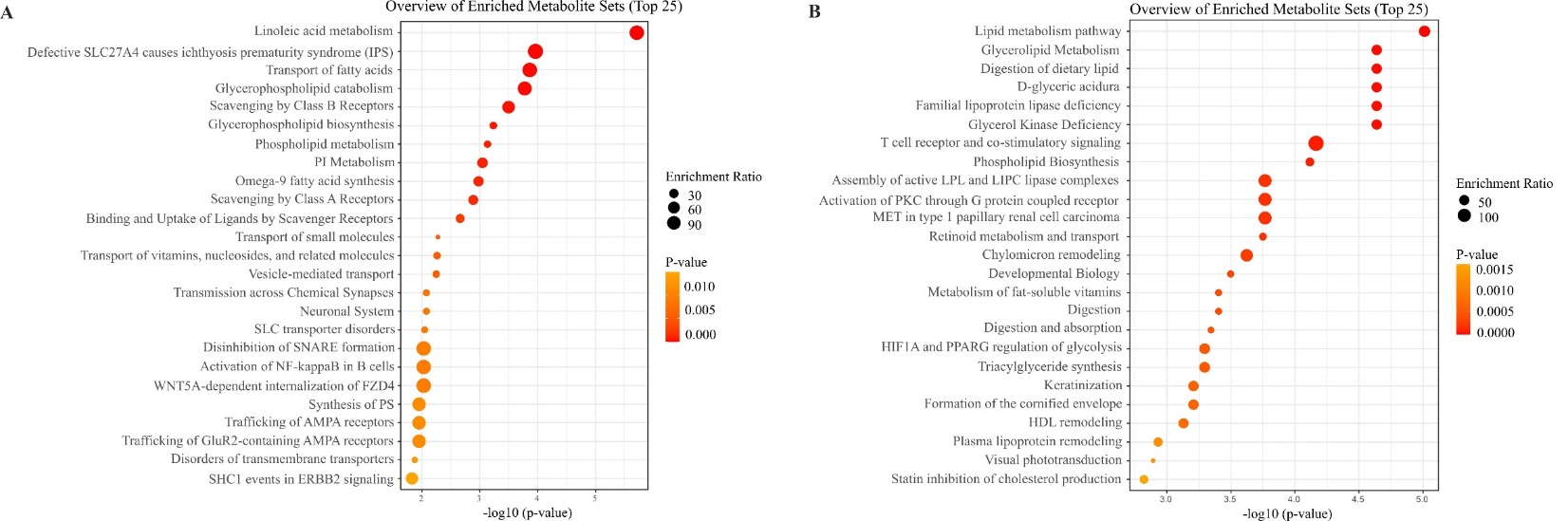

**Figure S3**. Enrichment analysis of different lipophilic metabolites in JIMT1 and SKBR3. (A) Metabolic pathway enrichment of lipophilic metabolites upregulated in JIMT1. (B) Metabolic pathway enrichment of lipophilic metabolites downregulated in JIMT1.

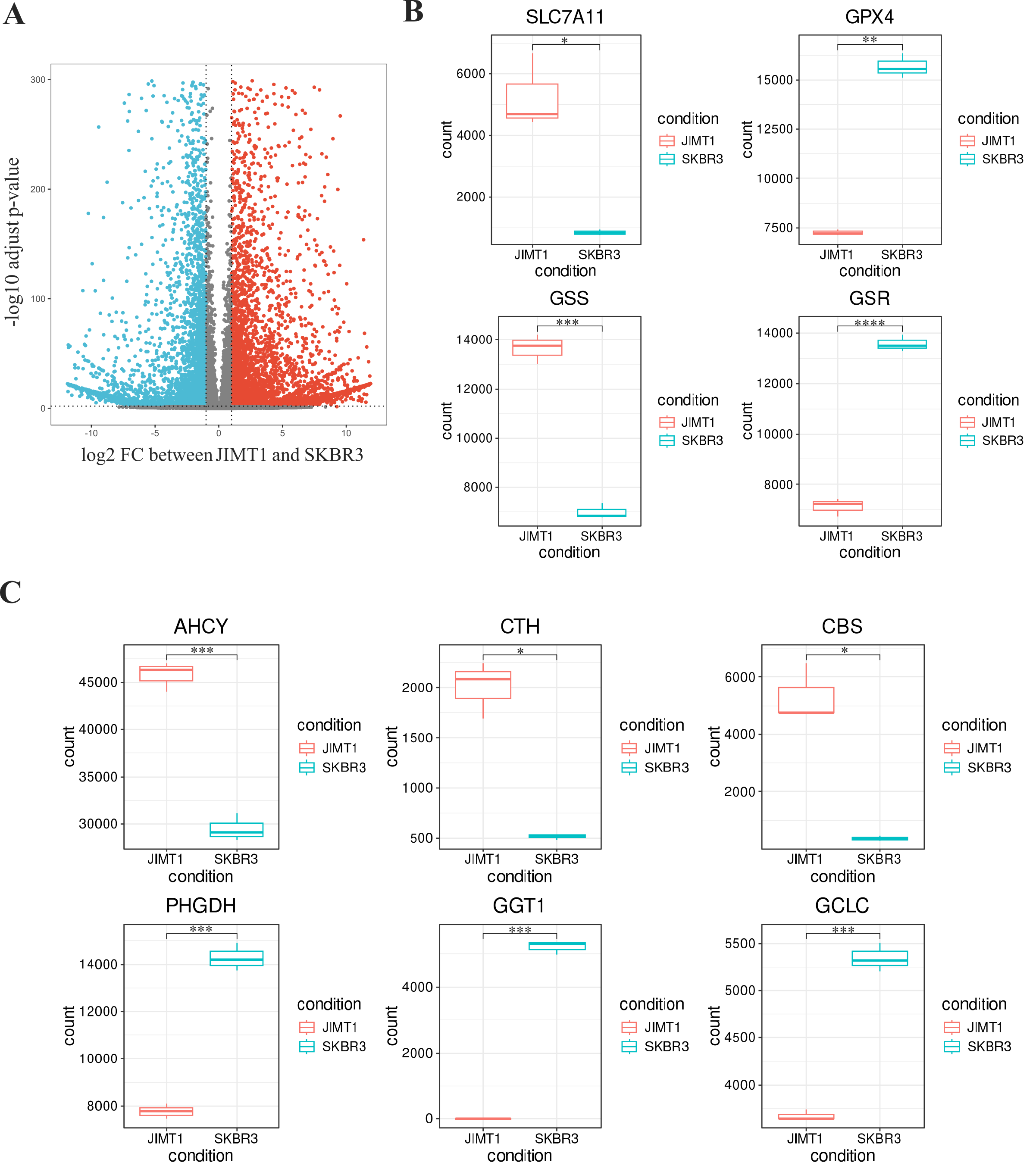

**Figure S4.** Analysis of transcriptomic data in JIMT1 and SKBR3. (A) Volcano plot of different genes in JIMT1 and SKBR3. (B) Transcriptional expression of SLC7A11, GPX4, GSS and GSR in JIMT1 and SKBR3. (C) Transcriptional expression of key genes involved with transsulfuration activity and γ-glutamyl-peptides synthesis activity (AHCY, CTH, CBS, PHGDH, CGT1 and GCLC) in JIMT1 and SKBR3.

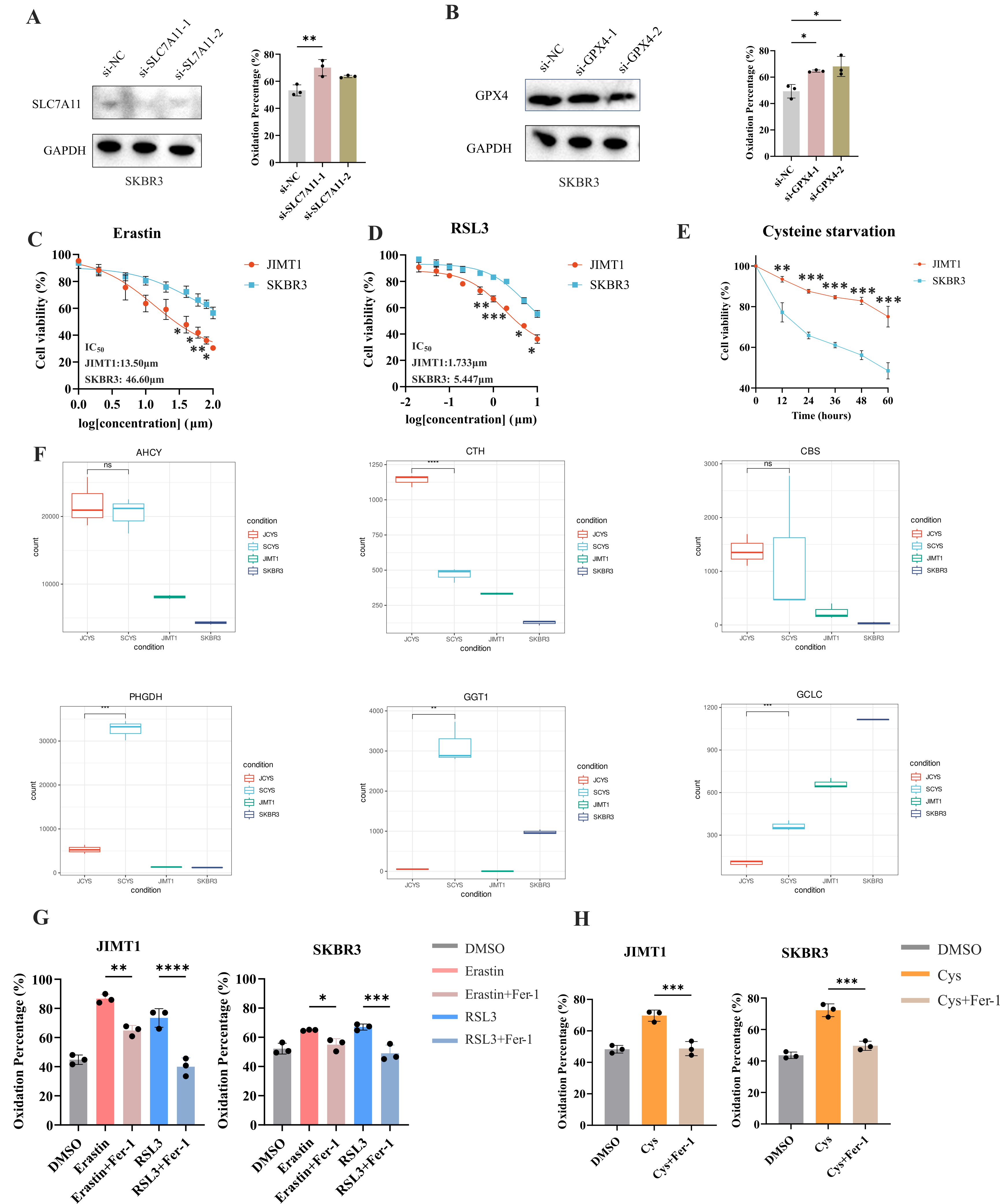

**Figure S5**. Knockdown of genes and targeting cysteine metabolism in JIMT1 and SKBR3. (A) Knockdown of SLC7A11 could increase lipid peroxidation in SKBR3. (B) Knockdown of GPX4 could increase lipid peroxidation in SKBR3. (C) Different sensitivity of JIMT1 and SKBR3 to treatment with erastin for 24 hours. (D) Different sensitivity of JIMT1 and SKBR3 to treatment with RSL3 for 24 hours. (E) Different sensitivity of JIMT1 and SKBR3 to treatment with cysteine starvation. (F) Transcriptional expression of key genes involved with transsulfuration activity and γ-glutamyl-peptides synthesis activity (AHCY, CTH, CBS, PHGDH, CGT1 and GCLC) in JIMT1 and SKBR3 after cysteine starvation. JCYS, JIMT1 cysteine starvation. SCYS, SKBR3 cysteine starvation. (G) The utilization of Fer-1 (10μm) inhibits lipid peroxidation resulted from erastin and RSL3. (H) The utilization of Fer-1 (10μm) inhibits lipid peroxidation resulted from cysteine starvation. Cys, cysteine starvation.

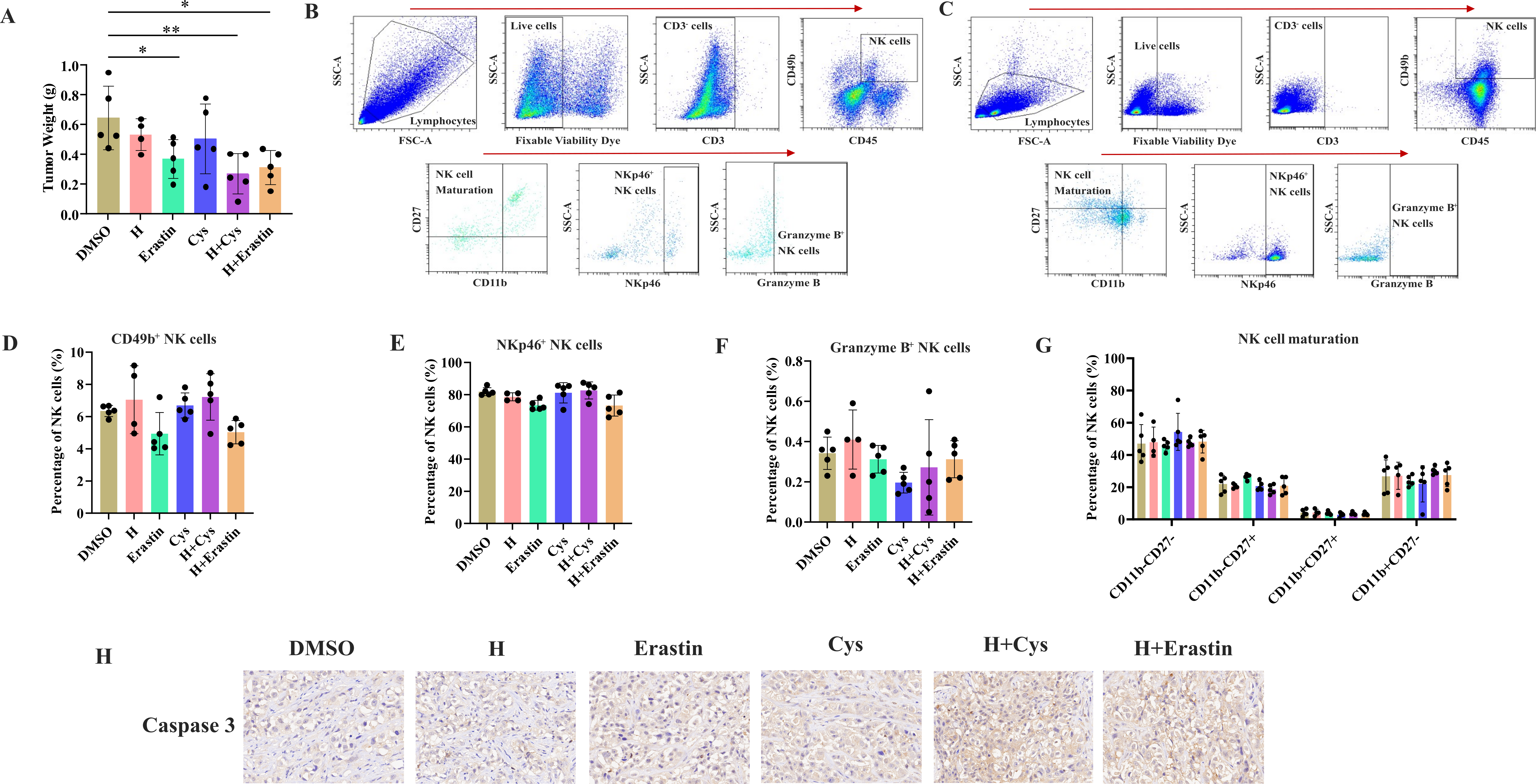

**Figure S6**. Detection of immune cell compositions in tumor and spleen samples. (A) Tumor weight of different treatment groups. (B-C) Strategies in the flow cytometry analysis of NK cells in tumor (B) and spleen (C). (D-G) Abundance of CD49b+ (D), NKp46+ (E), Granzyme B+ (F) and different development status (G) NK cells in spleen. (H) Representing immunohistochemistry images caspase 3 in different treatment groups.

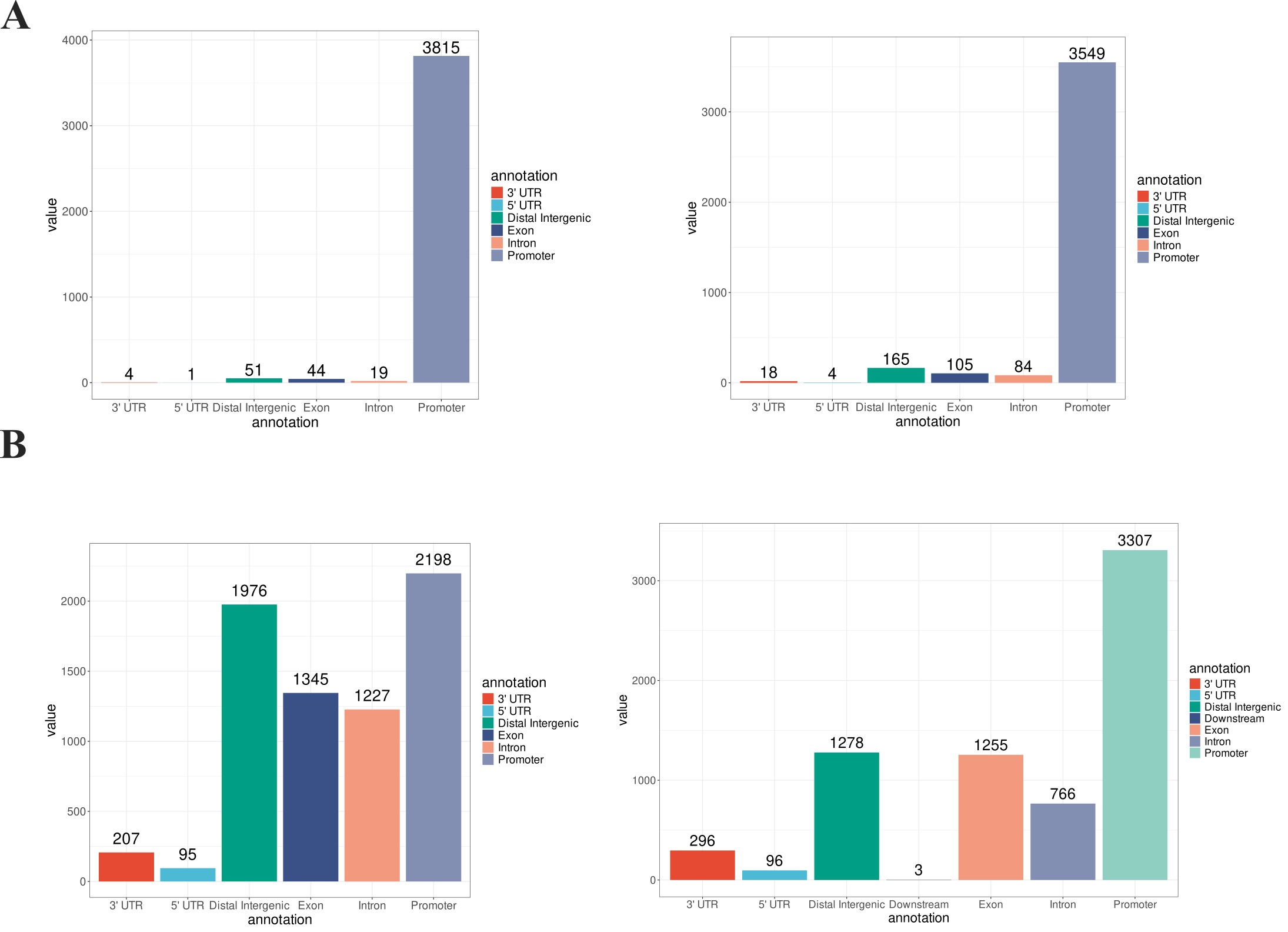

**Figure S7.** Location of altered H3K4me3 and H3K27me3 peaks in JIMT1 and SKBR3. (A) Enriched H3K4me3 peaks in JIMT1 (left) and SKBR3 (right). (B) Enriched H3K27me3 peaks in JIMT1 (left) and SKBR3 (right).

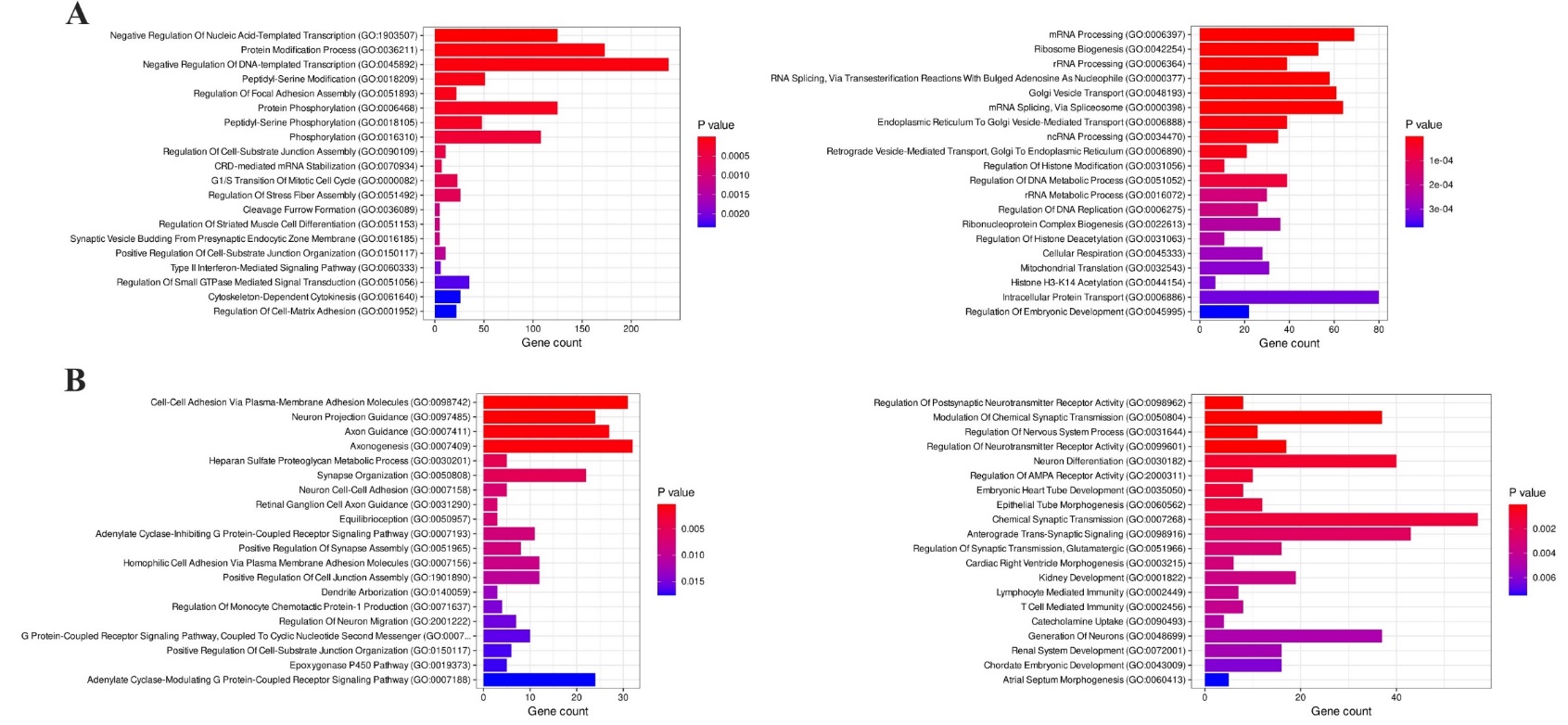

**Figure S8.** Enrichment analysis of biological processes related to H3K4me3 and H3K27me3 alterations. (A) Biological processes involved with upregulated H3K4me3 in JIMT1 (left) and SKBR3 (right). (A) Biological processes involved with upregulated H3K27me3 in JIMT1 (left) and SKBR3 (right).

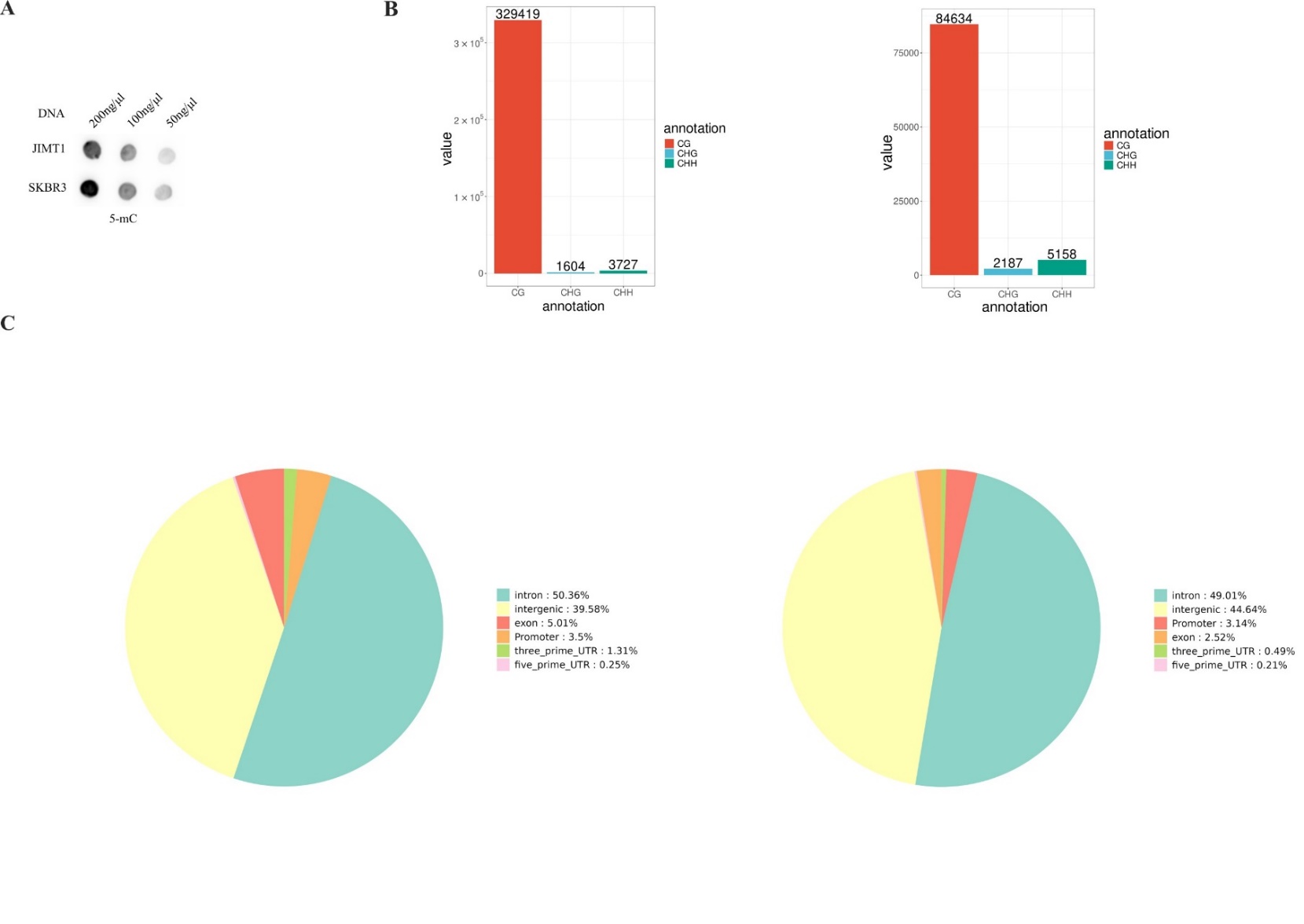

**Figure S9**. Different DNA methylation status between JIMT1 and SKBR3. (A) Dot blot of 5-mC levels in JIMT1 and SKBR3. (B) Enriched differentially methylated regions (DMR) in JIMT1 (left) and SKBR3 (right). (C) Location of CG-DMRs in JIMT1 (left) and SKBR3 (right) across the whole genome.

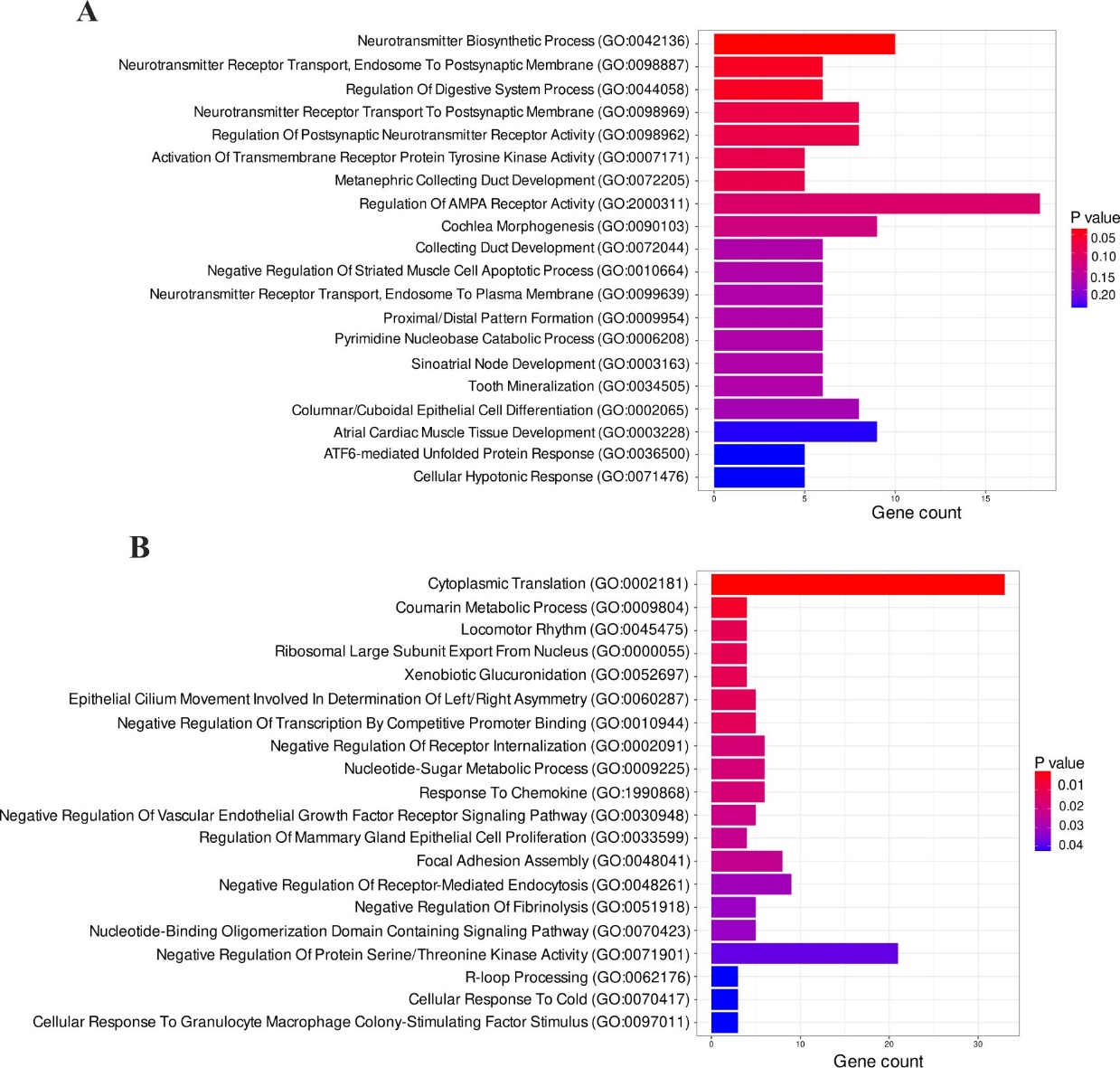

**Figure S10**. Enrichment analysis of biological processes related to DNA methylation alterations. (A) Biological processes associated with upregulated DNA methylation in JIMT1. (B) Biological processes associated with upregulated DNA methylation in SKBR3.

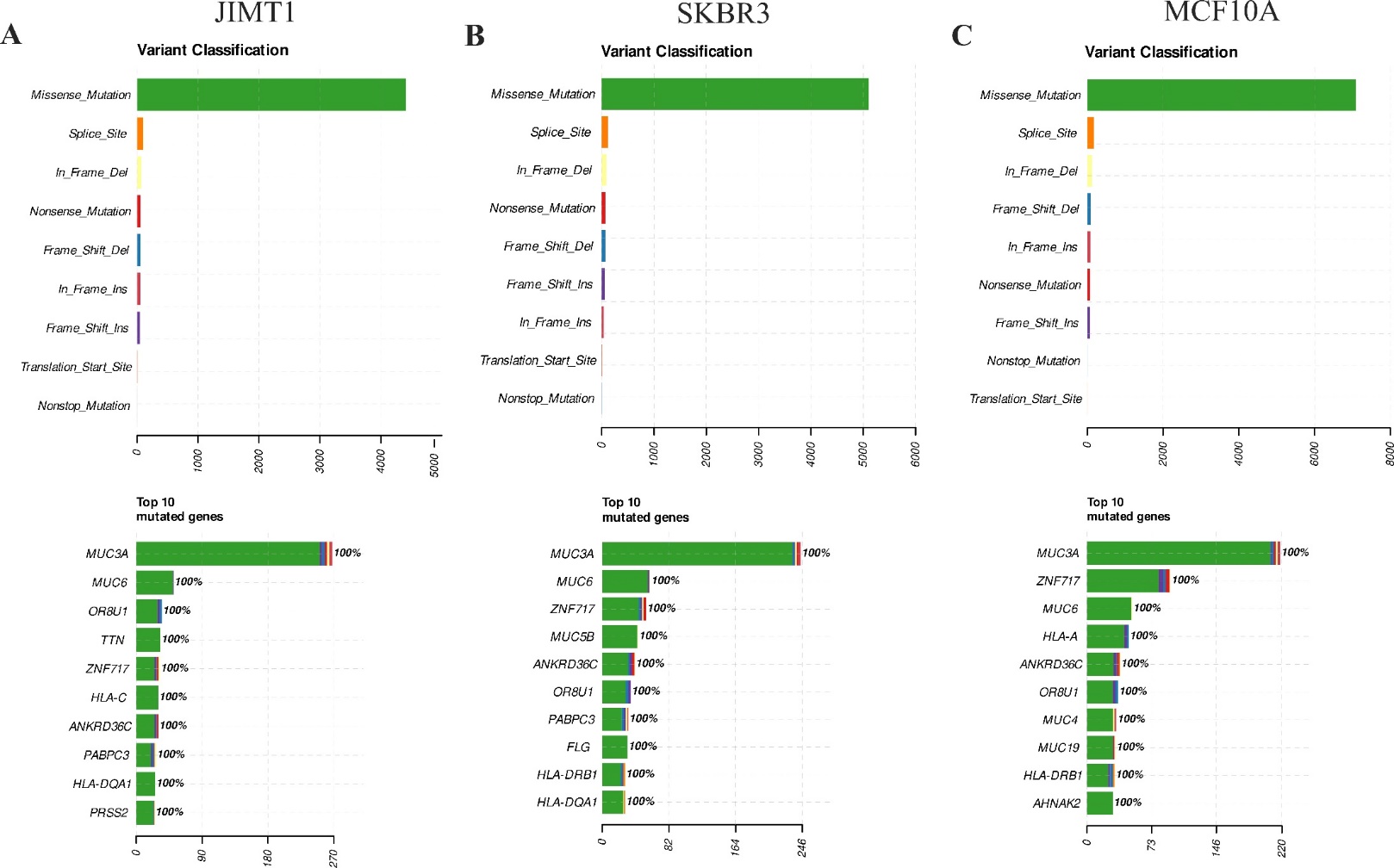

**Figure S11**. Genomic variations in JIMT1, SKBR3 and MCF10A. (A) Variant classification and top 10 mutated genes in JIMT1 across the whole genome. (B) Variant classification and top 10 mutated genes in SKBR3 across the whole genome. (C) Variant classification and top 10 mutated genes in MCF10A across the whole genome.

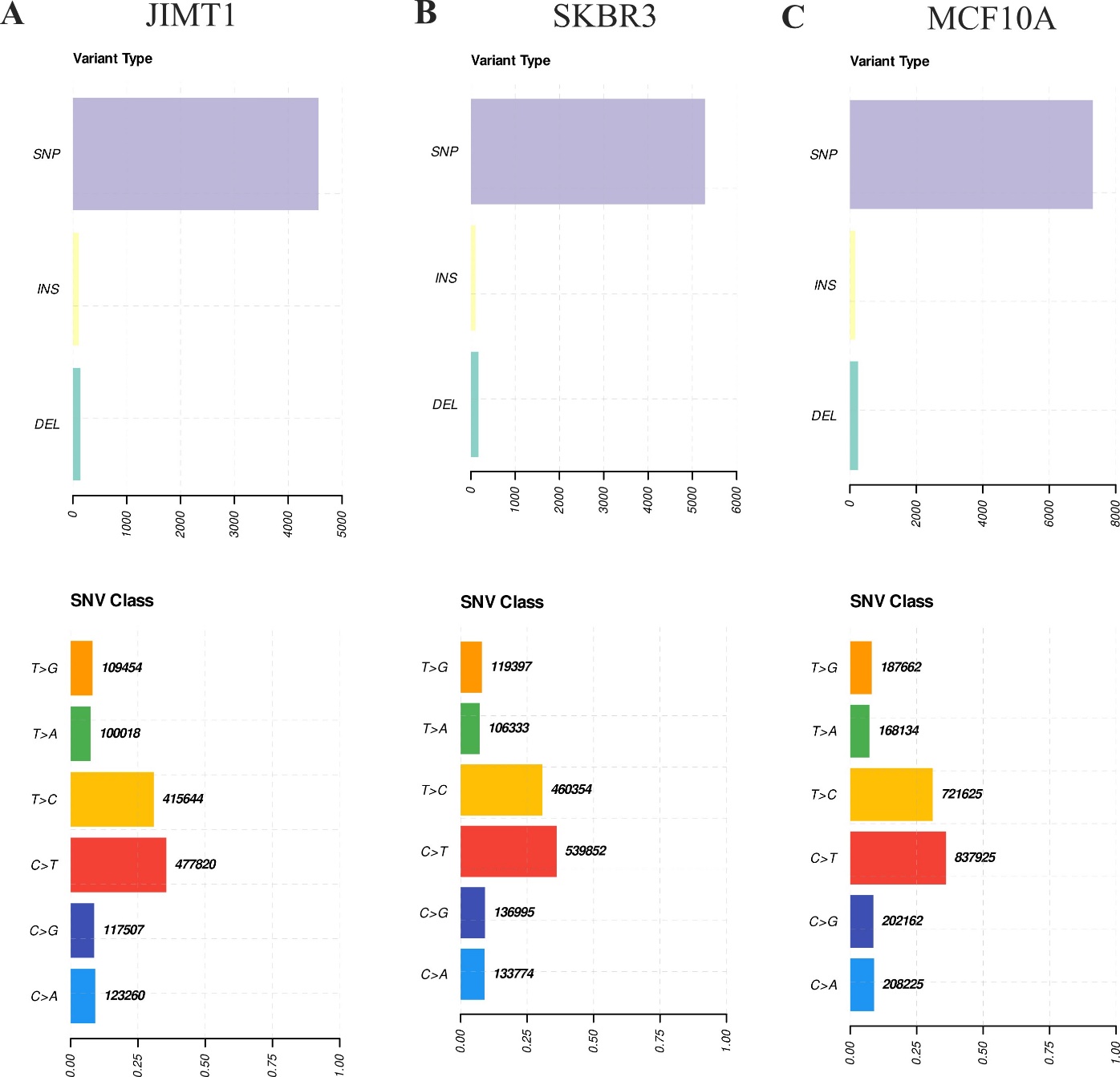

**Figure S12**. SNV characteristics in JIMT1, SKBR3 and MCF10A. (A) The most frequent SNV type in JIMT1. (B) The most frequent SNV type in SKBR3. (C) The most frequent SNV type in MCF10A. SNV, single nucleotide variation.

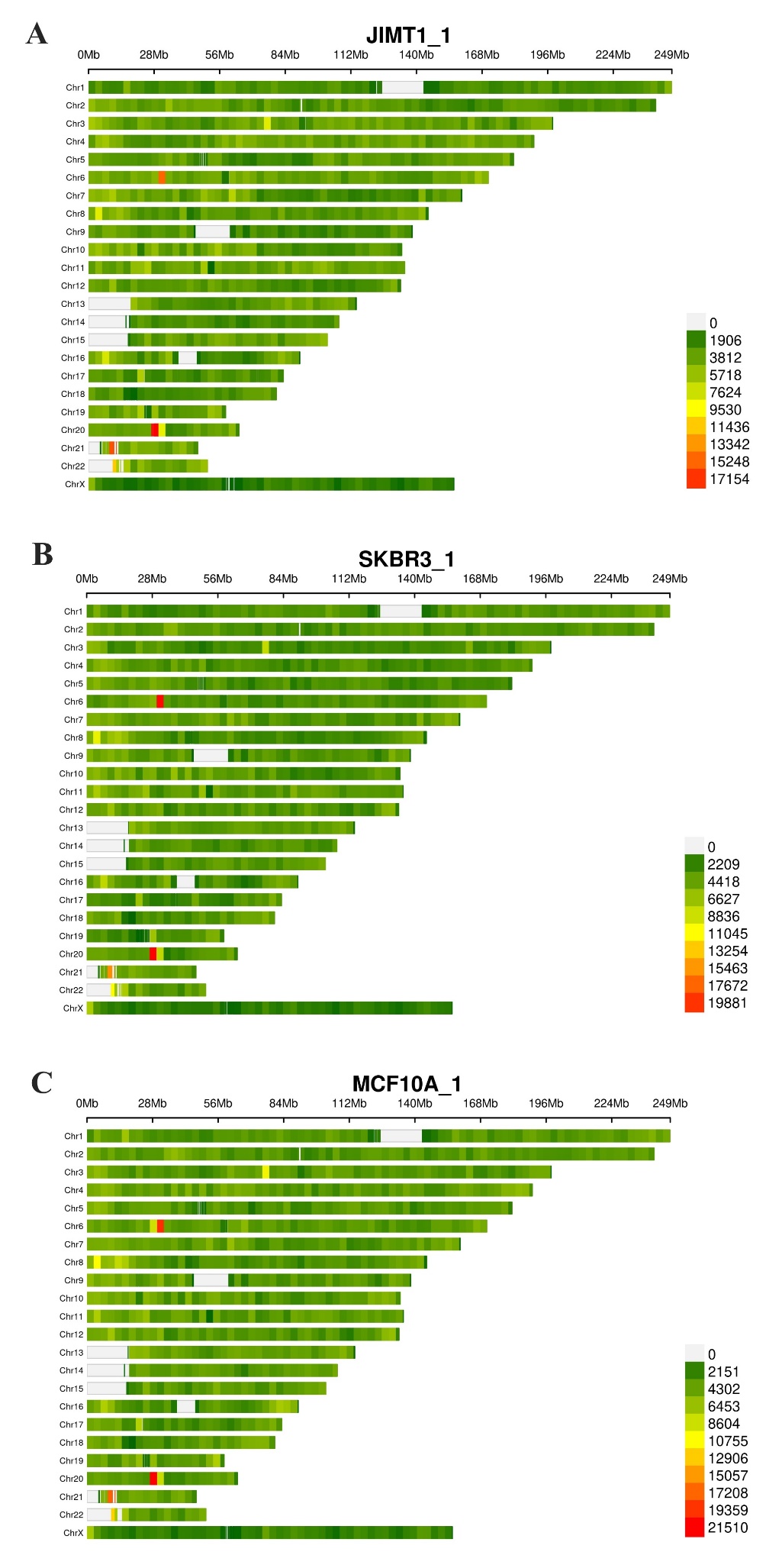

**Figure S13**. Density of SNP in JIMT1 (A), SKBR3 (B) and MCF10A (C) across the whole genome. SNP, single nucleotide polymorphism.

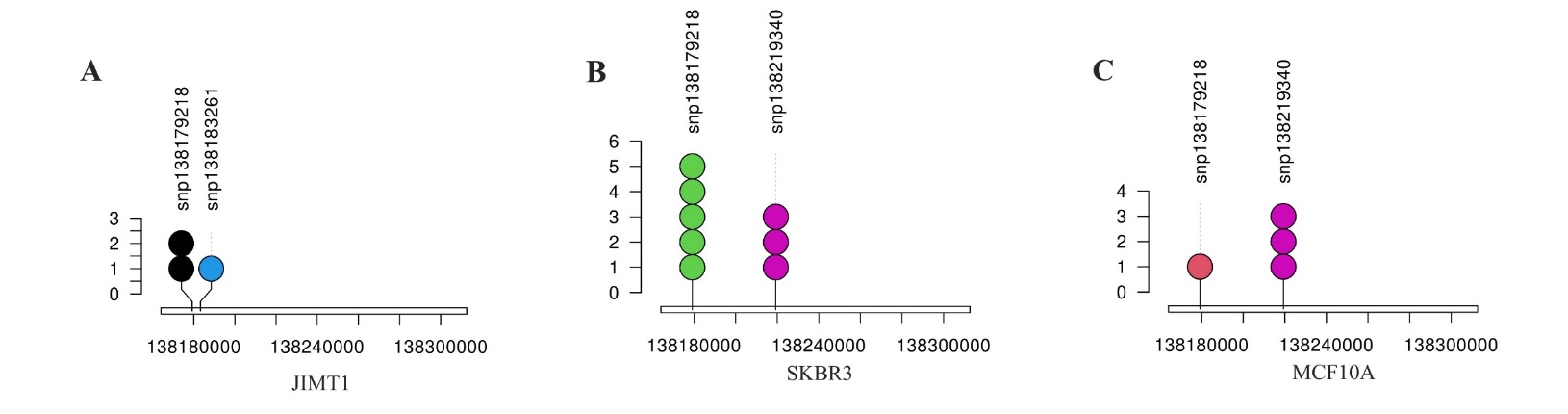

**Figure S14**. SNVs located in the exon region of SLC7A11 in JIMT1 (A), SKBR3 (B) and MCF10A (C).

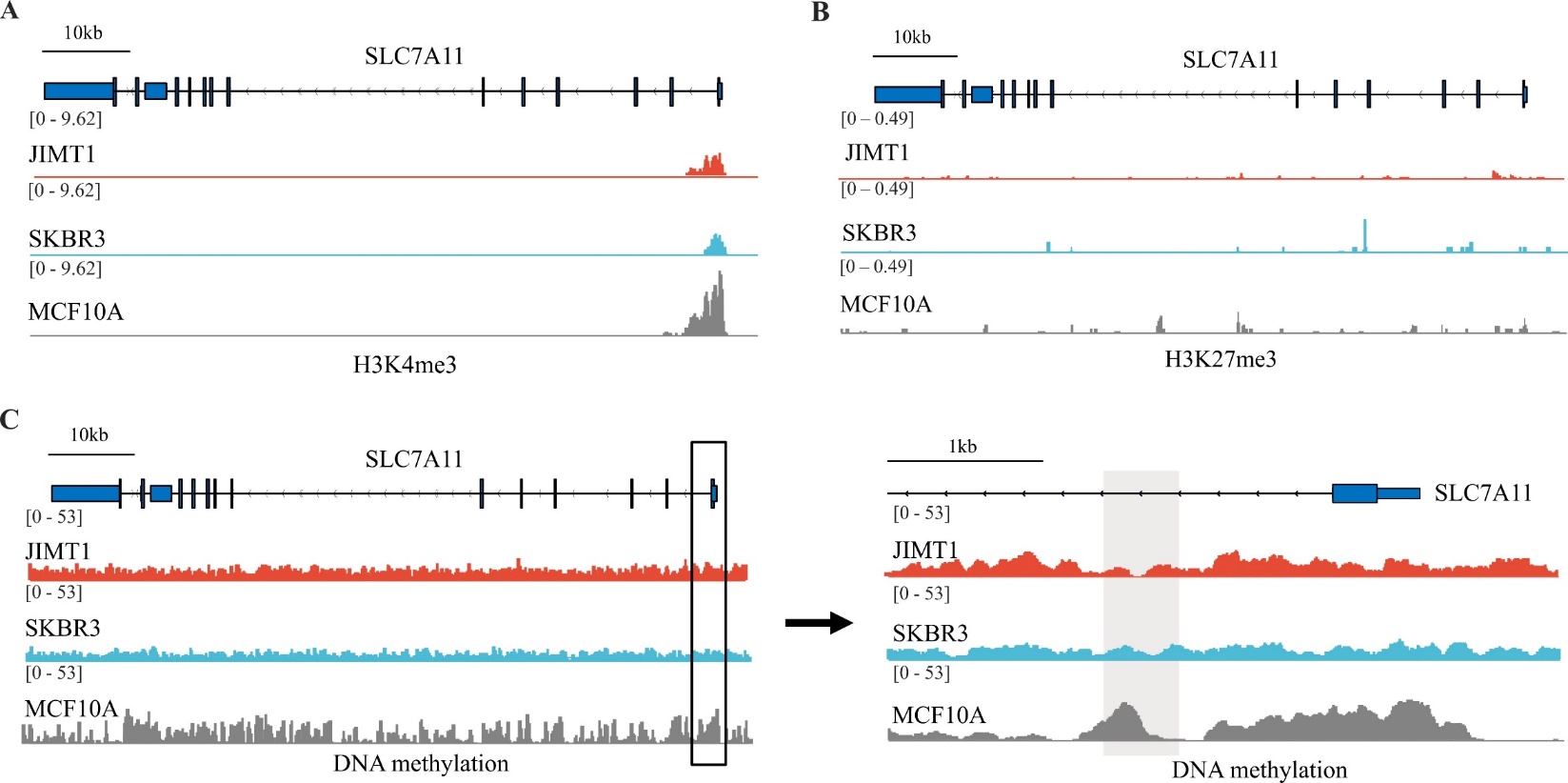

**Figure S15**. Different histone modifications and DNA methylation in SLC7A11 between JIMT1, SKBR3 and MCF10A. (A-B) H3K4me3 (A) and H3K27me3 (B) peaks located in SLC7A11. (C) Abundance of 5-mC at SLC7A11 promoter regions.

| **siRNA** | **5'-3' sequence** | |
| --- | --- | --- |
| si-ASH2L-1 | sense strand | CGAAGACAAUGUUCUCCAA(dT)(dT) |
|  | antisense strand | UUGGAGAACAUUGUCUUCG(dT)(dT) |
| si-ASH2L-2 | sense strand | GCUGACACAUUUGGCAUAGAU |
|  | antisense strand | AUCUAUGCCAAAUGUGUCAGC |
| si-SLC7A11-1 | sense strand | GGAGUUAUGCAGCUAAUUA(dT)(dT) |
|  | antisense strand | UAAUUAGCUGCAUAACUCC(dT)(dT) |
| si-SLC7A11-2 | sense strand | CUACUUUACGACCAUUAAU(dT)(dT) |
|  | antisense strand | AUUAAUGGUCGUAAAGUAG(dT)(dT) |
| si-GPX4-1 | sense strand | GGAGUAACGAAGAGAUCAA(dT)(dT) |
|  | antisense strand | UUGAUCUCUUCGUUACUCC(dT)(dT) |
| si-GPX4-2 | sense strand | GGAAGUGGAUGAAGAUCCA(dT)(dT) |
|  | antisense strand | UGGAUCUUCAUCCACUUCC(dT)(dT) |

**Table S1. Oligonucleotides sequences of siRNAs.**

| **sgRNA** | **5'-3' sequence** |
| --- | --- |
| SLC7A11-sgRNA-1 | (mU)*(mA)*(mC)*GAAAAAUAAGCCCAACGGUUUUAGAGCUAGAAAUAGCAAGUUAAAAUAAGGCUAGUCCGUUAUCAACUUGAAAAAGUGGCACCGAGUCGGUGCU*(mU)*(mU)*(mU) |
| SLC7A11-sgRNA-2 | (mA)*(mU)*(mG)*AAUUGAUUGGUACACAUGUUUUAGAGCUAGAAAUAGCAAGUUAAAAUAAGGCUAGUCCGUUAUCAACUUGAAAAAGUGGCACCGAGUCGGUGCU*(mU)*(mU)*(mU) |

**Table S2. Oligonucleotides sequences of sgRNAs.**

| **PCR primers** | **5'-3' sequence** | |
| --- | --- | --- |
| SLC7A11-ChIP | Forward | CTCAGCTTCCTCATGGGCTT |
|  | Reverse | GCTGAGTAATGCTGGAGGCT |
| SLC7A11-MeDIP | Forward | CTCCTGTTCCCATGTGTACCAA |
|  | Reserve | AGTTCTCTAGATGAGTGTCAAACCA |

**Table S3. PCR primers sequences for ChIP and MeDIP tests.**
